# Exposure to cooler temperatures during pupal development increases *Aedes aegypti* vector competence and the *R*_*0*_ for Zika virus

**DOI:** 10.1101/2024.02.16.580619

**Authors:** Tyler D. Pohlenz, Jeremy Vela, Martin Reyna, Chris Fredregill, Byul Hyur, Madhav Erraguntla, Mark Lawley, Mustapha Debboun, Zach N. Adelman, Martial L. Ndeffo-Mbah, Kevin M. Myles

## Abstract

Temperature profoundly affects various aspects of ectotherm biology. Notably, in mosquito species that spread viral diseases, temperature influences not only vector biology, but also the dynamics of pathogen-vector interactions. However, research attempting to address the role of the thermal environment in disease transmission often employs constant temperatures, which do not reflect the natural diurnal fluctuations these organisms experience. Additionally, most studies focus on adult mosquitoes in the period following virus infection. Much less attention has been paid to evidence indicating that temperatures experienced during earlier developmental stages may also affect the ability of disease vectors to be infected with and transmit viruses. Here, we show that *Aedes aegypti* exposed to temperatures below 25°C, specifically during the pupal stage of development, exhibit heightened susceptibility to Zika virus (ZIKV), which increases transmission efficiency. Modeling suggests that exposing mosquitoes to cooler fluctuating diurnal temperature ranges only during the relatively short pupal stage increases the R_0_ or reproductive number of ZIKV. Data loggers placed near Harris County Mosquito Control trap sites consistently recorded temperatures below 25°C, indicating natural exposure to such conditions. These results highlight the significance of thermal heterogeneity in the microhabitats where container-breeding mosquitoes undergo development. Such heterogeneity may play a more important role in the transmission of mosquito-borne diseases than previously recognized.

**Author Summary:** A paucity of information regarding how natural heterogeneity in thermal environments influences the spread of viral pathogens by mosquito species hinders our ability to decipher current and future patterns of disease transmission under changing climatic conditions. Here, we show that *Ae. aegypti* pupae exposed to cooler fluctuating diurnal temperature ranges, mimicking realistic field conditions, increases vector competence for ZIKV. Modeling cooler temperatures during immature life stages predicts slower development and increased mortality, counterbalancing increases in disease transmission. However, exposing mosquitoes to lower temperatures only during the relatively short pupal stage was predicted to increase the R_0_ value of ZIKV. These results suggest that environmental temperatures specifically experienced during the pupal stage may have an important role in disease transmission.

## Introduction

Understanding the pivotal role of temperature in the transmission of arthropod-borne viruses (arboviruses) is of significant importance. This knowledge is crucial for deciphering the current spatiotemporal patterns of disease transmission, and forecasting future epidemic disease dynamics amidst rapidly changing climatic conditions (1–3). However, modeling and predicting alterations in the disease dynamics of mosquito-borne viral pathogens presents formidable challenges due to the intricate interplay of environmental factors on both viruses and vectors (4–7). While numerous models of arboviral disease transmission incorporate the impact of environmental temperatures on parameters like mosquito lifespan (8, 9), duration of the developmental cycle (10, 11), and extrinsic incubation period of a specific arbovirus post-exposure (12–16), our grasp of how natural variability in thermal environments influences disease transmission dynamics remains incomplete. Further, studies have revealed that the temperatures experienced by mosquitoes prior to virus exposure can also influence vector competence (17–21). Lower temperatures, in particular, have been observed to adversely affect a vector’s ability to modulate viral infections. Notably, there seems to exist an inverse relationship between the temperature at which mosquitoes undergo development and their subsequent susceptibility to virus infections as adults. These findings can be correlated with the microclimate temperatures measured within mosquito breeding sites during periods when vector-borne disease transmission has historically been characterized as high. For instance, during a dengue virus (DENV) transmission season in Buenos Aires (January-March), temperatures in shaded microhabitats were estimated to be approximately 10°C cooler, ranging from 22-25°C, in contrast with sunlit areas where temperatures ranged between 30-37°C (22). Similarly, during an outbreak of chikungunya virus (CHIKV) in La Réunion, *Aedes albopictus* mosquitoes were observed to prefer shaded breeding sites with average temperatures as low as 12.6°C (23).

The body temperatures of small ectothermic animals such as mosquitoes are significantly influenced by the constantly changing microclimatic conditions of their surroundings (1, 24). Biophysical models suggest that the body temperatures of ectotherms quickly equilibrate to match the ambient temperature. Consequently, the thermal characteristics and physical attributes of the environments where these organisms develop play a critical role in determining their body temperatures, especially for arthropods. However, most research studies investigating the impact of temperature on the biology of disease vectors and arboviruses typically employ constant temperature settings, which may not accurately represent the natural temperature fluctuations experienced by these organisms (21, 25). Further, we are only aware of a single study that has explored the effects of daily temperature fluctuations before virus exposure on the transmission of pathogens. In this particular study (26), *Ae. albopictus* mosquitoes were raised outdoors during either summer (August-September) or autumn (September-November) months, and then exposed to DENV. Those raised in the cooler autumn months showed higher transmission potential than did those from the warmer summer months. However, these experiments did not attempt to pinpoint the specific life stage of the mosquito responsible for the observed temperature dependent effects on vector competence.

In this study, we demonstrate that *Aedes aegypti* mosquitoes subjected to temperatures ≤ 25°C, particularly during their pupal development phase, are more susceptible to Zika virus (ZIKV) infection resulting in a higher transmission efficiency. Exposing immature mosquito stages to cooler temperatures can result in significant delays in development time and increase mortality rates. These factors would normally offset any observed increases in vector competence when modeling the pathogen’s basic reproductive number (*R*_0_). However, our simulations indicate that subjecting mosquitoes to cooler temperatures exclusively during the short pupal phase (1-3 days) can increase the R_0_ of ZIKV. Temperature monitors located at mosquito control trap sites in Harris County consistently registered temperatures under 25°C, showing that mosquitoes naturally experience such conditions. These results suggest that thermal heterogeneity in the microhabitats where container-breeding mosquitoes undergo development may have a more important role in the transmission of mosquito-borne diseases than previously recognized.

## Results

### Vector competence of Ae. aegypti reared under different constant temperatures

In accordance with the experimental design outlined in Figure 1, we investigated how different rearing temperatures affected the ability of *Ae. aegypti* to transmit ZIKV. To assess whether cooler environmental conditions make mosquitoes more susceptible to ZIKV, we reared Liverpool strain *Ae. aegypti* at constant temperatures ranging from 18°C to 25°C and provided them with blood meals containing equivalent virus titers (1 x 10^6^ pfu/ml). Our control group was raised at a constant temperature of 28°C. Fourteen days after consuming the blood meal, we counted the number of ZIKV-infected mosquitoes and compared it with the control group. We found that mosquitoes reared at ≤ 25°C were more likely to be infected with ZIKV in comparison to the control group, despite all groups being kept at 28°C during the virus incubation period (Fig. 2A). We also analyzed the heads of ZIKV-infected mosquitoes for evidence of systemic infection. Rearing mosquitoes at constant temperatures of ≤ 25°C significantly enhanced the dissemination of ZIKV from the midguts of infected mosquitoes (Fig. 2B). Real-time quantitative PCR (RT-qPCR) analysis on infected mosquitoes revealed that the increased susceptibility and dissemination observed in mosquitoes reared at lower temperatures correlated with significantly higher levels of viral replication (Fig. 2C). These findings are consistent with our previous research on this topic, which demonstrated that mosquitoes reared at a constant temperature of 18°C before consuming infectious blood meals containing either CHIKV or yellow fever virus (YFV) exhibited increases in infection prevalence and viral load (17). In the present study, virus transmission rates were also monitored during and after the extrinsic incubation period. There was no significant difference in transmission rates between mosquitoes reared at temperatures ≤25°C and the control mosquitoes raised at 28°C (Fig. S1A). However, due to the increased prevalence of midgut and systemic infections in mosquitoes reared at lower temperatures, transmission efficiency increased significantly (Fig. 2D). Transmission efficiency here refers to the percentage of mosquitoes ingesting ZIKV that could subsequently transmit the pathogen.

**Fig. 1.**
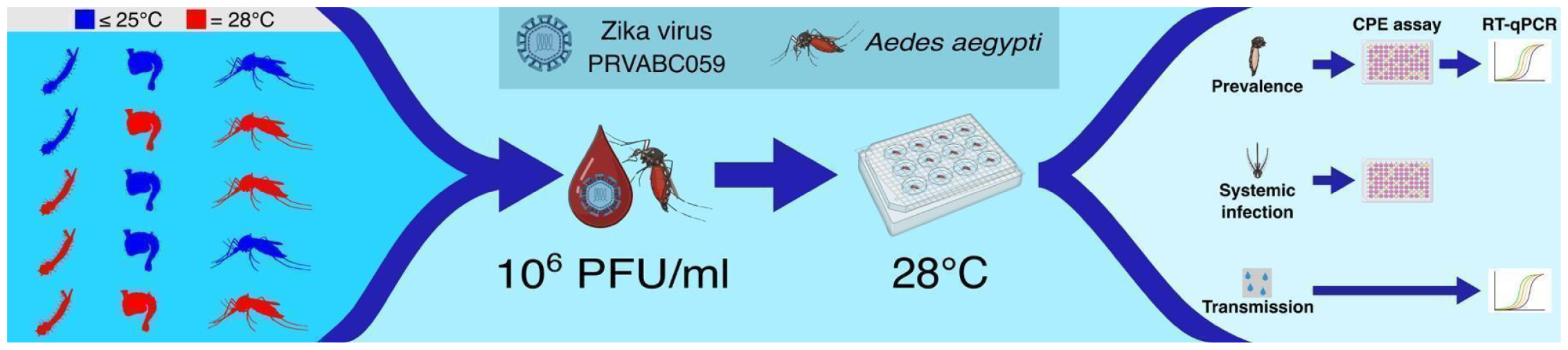
Experimental design of vector competence studies. *Ae. aegypti* (Liverpool strain) mosquitoes were reared at different temperatures, either for their entire lifespan or only during specific life stages. Three to five day old mosquitoes were exposed to a blood meal containing 1 x 10^6^ PFU/ml of ZIKV PRVABC59. Subsequently, engorged mosquitoes were individually isolated in chambers and held at 28°C during the extrinsic incubation period. To normalize biting rates, which might be influenced by exposure to lower temperatures, all mosquitoes were shifted to 28°C five to six hours before blood feeding. Throughout the study, mosquitoes were sustained on filter papers soaked in 10% sucrose, facilitating saliva collection during feeding. RT-qPCR was employed to test the saliva samples for the presence of the virus. At 14 days post-blood meal, individual mosquito bodies, heads, and corresponding filter papers were collected and preserved at −80°C until processing. CPE assays were performed using homogenates from bodies and heads, and viral RNA was extracted from mosquito body homogenates. Viral load was quantified through RT-qPCR. Positive bodies indicated midgut infection, positive heads signified systemic infection, and positive filter papers (Ct < 35) indicated transmission of ZIKV.

**Fig. 2.**
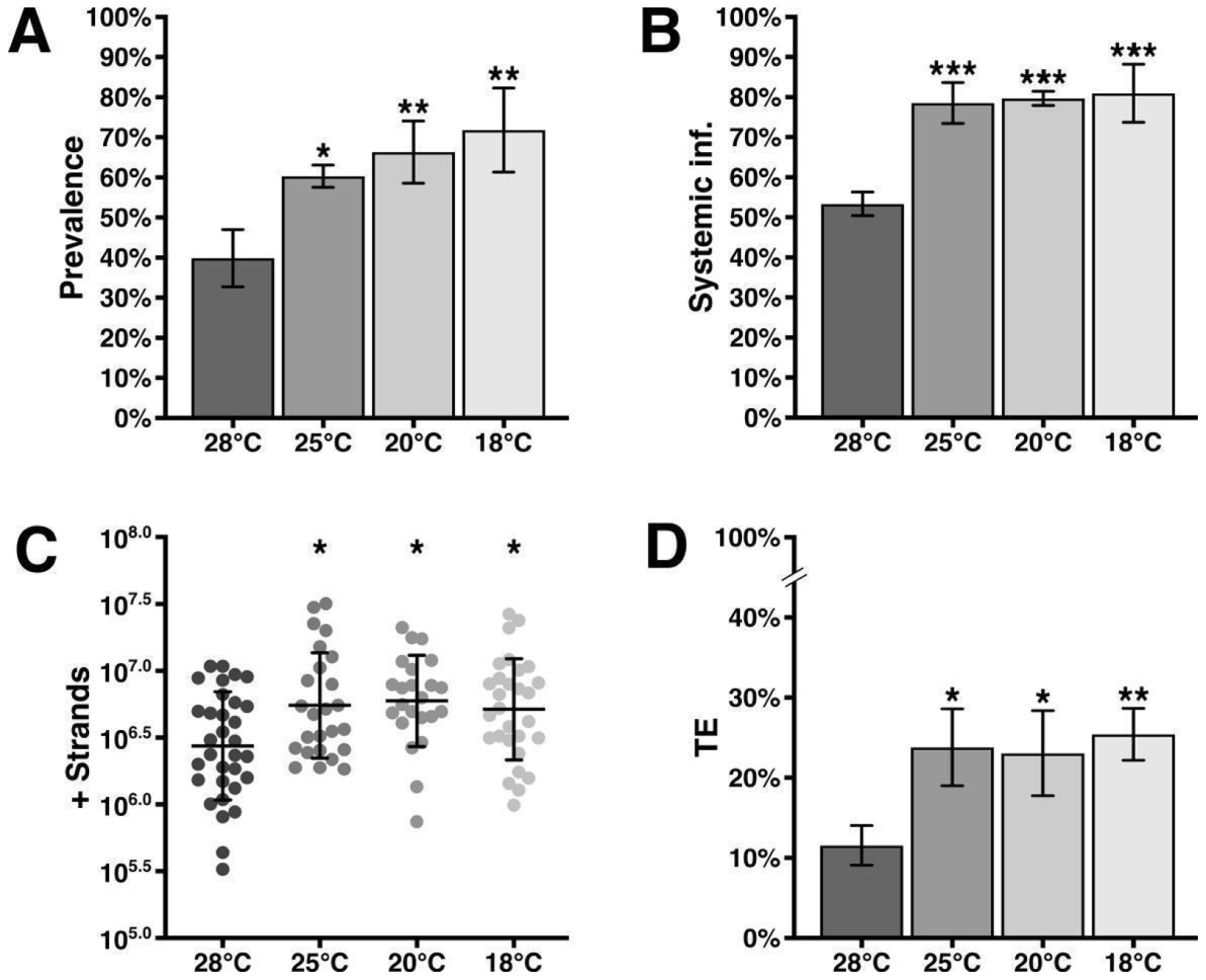
Vector competence of *Ae. aegypti* for ZIKV at various constant temperatures. (**A**) Prevalence of midgut infections; expressed as the proportion of positive bodies over the total number of mosquitoes that consumed a blood meal containing 1 x 10^6^ pfu/ml of ZIKV. (**B**) Prevalence of mosquitoes exhibiting systemic infections; expressed as the proportion of positive heads over positive bodies. (**C**) Quantification of viral genomes in ZIKV-infected mosquitoes; assessed using RT-qPCR. (**D**) Transmission efficiency; expressed as the proportion of ZIKV positive filter papers over the total number of mosquitoes that ingested the virus. All assessments were conducted 14 days post-blood meal and compared with a control group reared at a constant temperature of 28°C. Statistical significance was determined using one-way ANOVA with Dunnett’s multiple comparison test, where * indicates P < 0.05, ** indicates P < 0.01, and *** indicates P < 0.001.

### Vector competence of Ae. aegypti reared under heterogeneous temperatures

To investigate whether the previously observed increases in vector competence were linked to a specific pre-adult life stage, we conducted experiments where we exposed immature mosquitoes to heterogenous temperature conditions, as depicted in Figure 1. The control group was kept at a constant temperature of 28°C. These experiments revealed that it was only necessary to expose pupae to temperatures of 25°C or lower in order to observe a significant increase in the prevalence of ZIKV infections (Fig. 3A). Additionally, subjecting only the pupal stage to temperatures ≤ 25°C also led to a significant increase in the dissemination of ZIKV from the midguts of infected mosquitoes (Fig. 3B). The observed increases in virus susceptibility and dissemination were again accompanied by significant increases in viral replication levels (Fig. 3C). Interestingly, we did not detect any significant difference in the prevalence of ZIKV infections or viral load when only larval stages were exposed to 18°C, in comparison with the control group (Fig. 3A, B, and C). Consistent with the findings reported above, we once again did not observe any substantial difference in transmission rates when pupal stages were exposed to temperatures ≤ 25°C (see Fig. S1B). However, significant increases in transmission efficiency were again evident when only pupae were subjected to temperatures ≤ 25°C (see Fig. 3D).

**Fig. 3.**
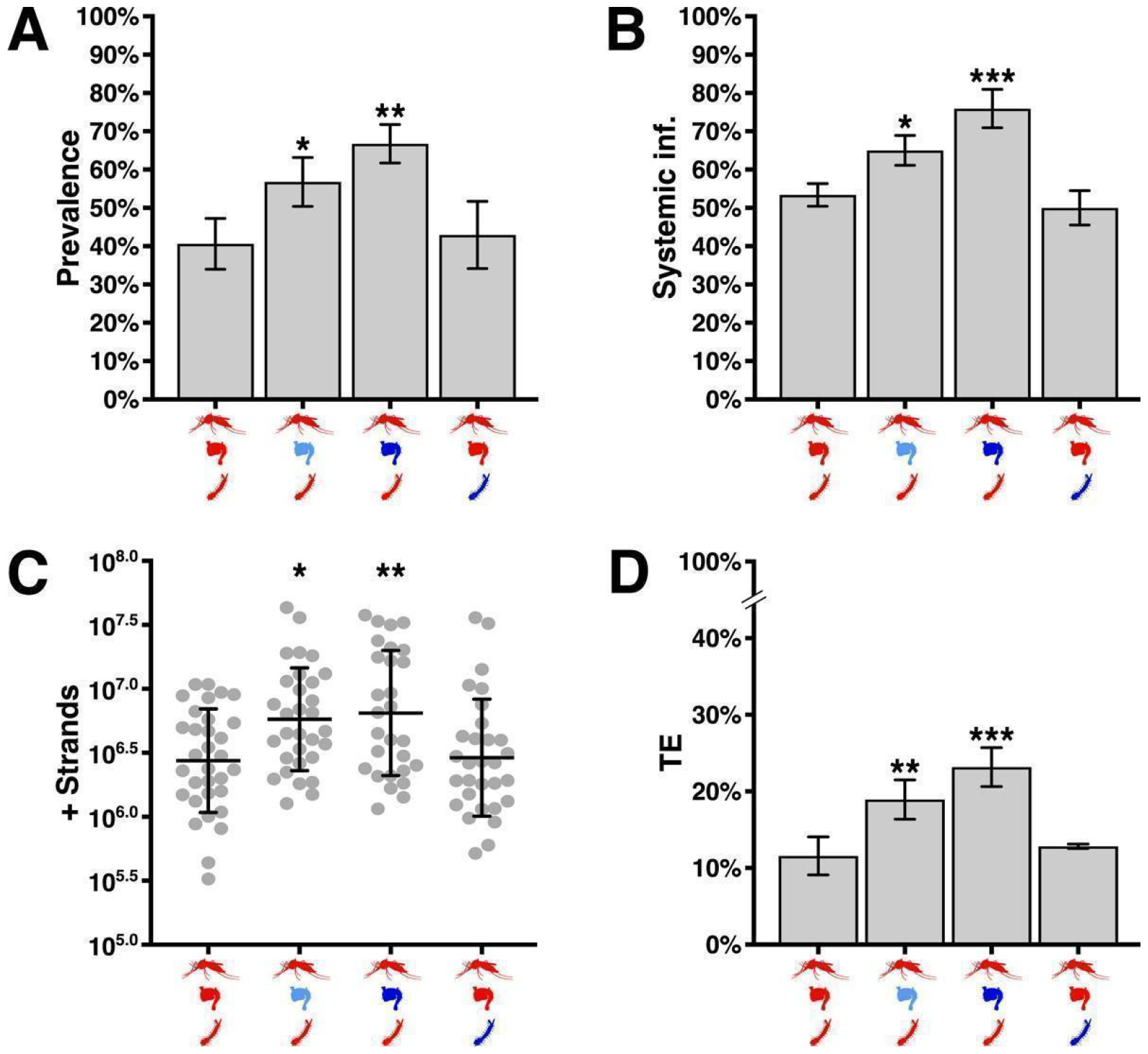
Vector competence of *Ae. aegypti* for ZIKV studied under heterogeneous temperature conditions. (**A**) Prevalence of midgut infections in mosquitoes exposed to different temperatures during specific life stages. Symbol colors represent the temperature at which each life stage was maintained, with red indicating 28°C, light blue indicating 25°C, and dark blue indicating 18°C. (**B**) Prevalence of mosquitoes exhibiting systemic infections. (**C**) Quantification of viral genomes in ZIKV-infected mosquitoes; assessed using RT-qPCR. (**D**) Transmission efficiency of ZIKV. All analyses were conducted 14 days post-blood meal (1 x 10^6^ PFU/ml ZIKV) and compared to a control group raised at a constant temperature of 28°C. Statistical significance was determined using one-way ANOVA with Dunnett’s multiple comparison test, where * indicates P < 0.05, ** indicates P < 0.01, and *** indicates P < 0.001.

### Vector competence of Ae. aegypti reared under selected diurnal temperature ranges

Unlike the stable temperatures often used in laboratory experiments, mosquitoes in their natural habitats experience fluctuating temperature patterns. To investigate this, we collaborated with Harris County Mosquito Control to install temperature sensors in different mosquito microenvironments located near trapping sites in the greater Houston, TX metropolitan area. Over the course of a year, we collected temperature data (Data S1) from these sensors. Using this data, we created five different diurnal temperature regimes (labeled Regimes I-V) that matched those recorded during specific time periods in the fall, a season often associated with higher levels of arbovirus transmission (see Fig. 4A). Regime I featured an average temperature of 29°C with a diurnal temperature range (DTR) of 7.8°C. Regime V, on the other hand, simulated an average temperature of 19°C with a DTR of 3.5°C. Cohorts reared under these two regimes served as high and low temperature control groups, respectively. Regimes II-IV replicated scenarios where temperatures dropped sharply (average of 19°C, DTR of 7.8°C), but only during the pupal development stage. In Regimes III and IV, recently emerged adult mosquitoes were also exposed to the low DTR before being administered infectious blood meals. With the exception of Regime IV, where adult mosquitoes continued to experience diurnal temperature fluctuations after exposure to the virus, all mosquitoes were moved to a constant temperature of 28°C after ingesting the infectious blood meal, in order to eliminate potential temperature-related effects on virus replication.

**Fig. 4.**
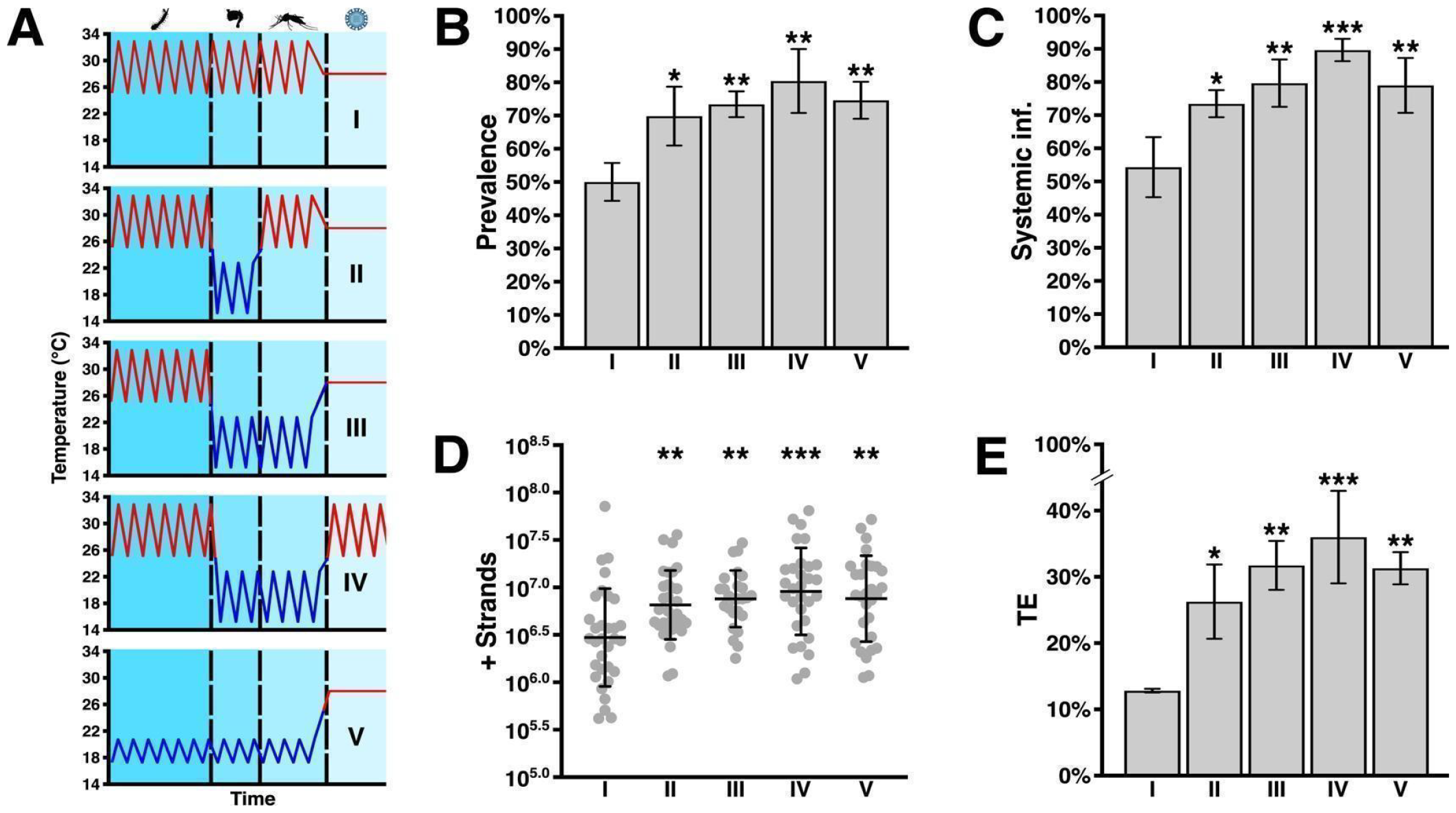
Vector competence of *Ae. aegypti* for ZIKV under fluctuating diurnal temperature conditions. (**A**) Temperature regimes mimicking natural diurnal temperature fluctuations. (**B**) Prevalence of midgut infections under the specified temperature regimes. (**C**) Prevalence of mosquitoes exhibiting systemic infections. (**D**) Quantification of viral genomes in ZIKV-infected mosquitoes; assessed using RT-qPCR. (**E**) Transmission efficiency of ZIKV. Mosquitoes were assessed 14 days post-blood meal (1 x 10^6^ PFU/ml ZIKV) and compared to the control group reared under temperature regime I. Statistical significance was determined using one-way ANOVA with Dunnett’s multiple comparison test, where * indicates P < 0.05, ** indicates P < 0.01, and *** indicates P < 0.001.

Consistent with our previous experiments, we again observed significant increases in the prevalence of midgut and systemic ZIKV infections (Fig. 4B and C), which also correlated with notably higher levels of viral replication (Fig. 4D), in mosquitoes reared under fluctuating diurnal temperatures ≤ 23°C. In comparison to the high DTR conditions (Regime I), there were generally no significant differences in transmission rates among mosquitoes kept at other diurnal temperature ranges. However, a noteworthy exception was observed in Regime IV, where mosquitoes continued to experience diurnal temperature fluctuations after virus exposure (Fig. S1C). In a previously published study (27), it was found that adult *Ae. aegypti* exposed to large diurnal temperature fluctuations with a low mean temperature (20°C) exhibited a significantly shorter extrinsic incubation period (EIP) when infected with DENV serotype 1. A shorter EIP suggests increased potential for virus transmission; although, such results have not been observed experimentally (21, 28–30). Intriguingly, *Ae. aegypti* exhibited higher rates of ZIKV transmission when exposed to the low mean temperature (19°C) and DTR over a relatively short duration of time, prior to introduction of the pathogen, after which they were returned to the DTR with a mean of 29°C. In any case, similar to our previous results, it was only necessary to expose the pupae to fluctuating diurnal temperatures ≤ 23°C in order to observe significant increases in transmission efficiency (Fig. 4E).

### Modeling R_0_ values for ZIKV

To further investigate how temperature-dependent changes in the vector competence of *Ae. aegypti* impact the transmission of ZIKV, we developed an extended version of a previously published mechanistic model for calculating the basic reproduction number (*R*_0_) of mosquito-borne viruses (4, 28, 31). Existing research has already highlighted the significant influence of temperature on critical parameters that are crucial for determining *R*_0_ values for arboviruses (1, 6, 28). What distinguishes our model is its explicit consideration of the effects of heterogeneous temperatures throughout both the juvenile and adult stages on ZIKV transmission—a previously unexplored aspect. We derived parameters for the *R*_0_ model from our experimental findings, as well as from prior empirical studies (4, 28), and proceeded to compute *R*_0_ values for ZIKV in comparison with mosquitoes reared at a constant 28°C. Lower temperatures were generally positively correlated with lower *R*_0_ values for ZIKV, when mosquitoes were reared at constant temperatures (Figure 5A); although, a minor increase in the *R*_0_ value of ZIKV was predicted at a constant temperature of 25°C. However, if only the pupal stage was subjected to cooler temperatures, a negative inverse relationship was observed between thermal environment and *R*_0_ values, with the most significant increase in the *R*_0_ value of ZIKV occurring at the lowest temperature (Figure 5B). This phenomenon is attributable to the effect that cooler temperatures have on the vector competence of adults when exposure occurs over the relatively short duration of time that mosquitoes spend in the pupal stage.

**Fig. 5.**
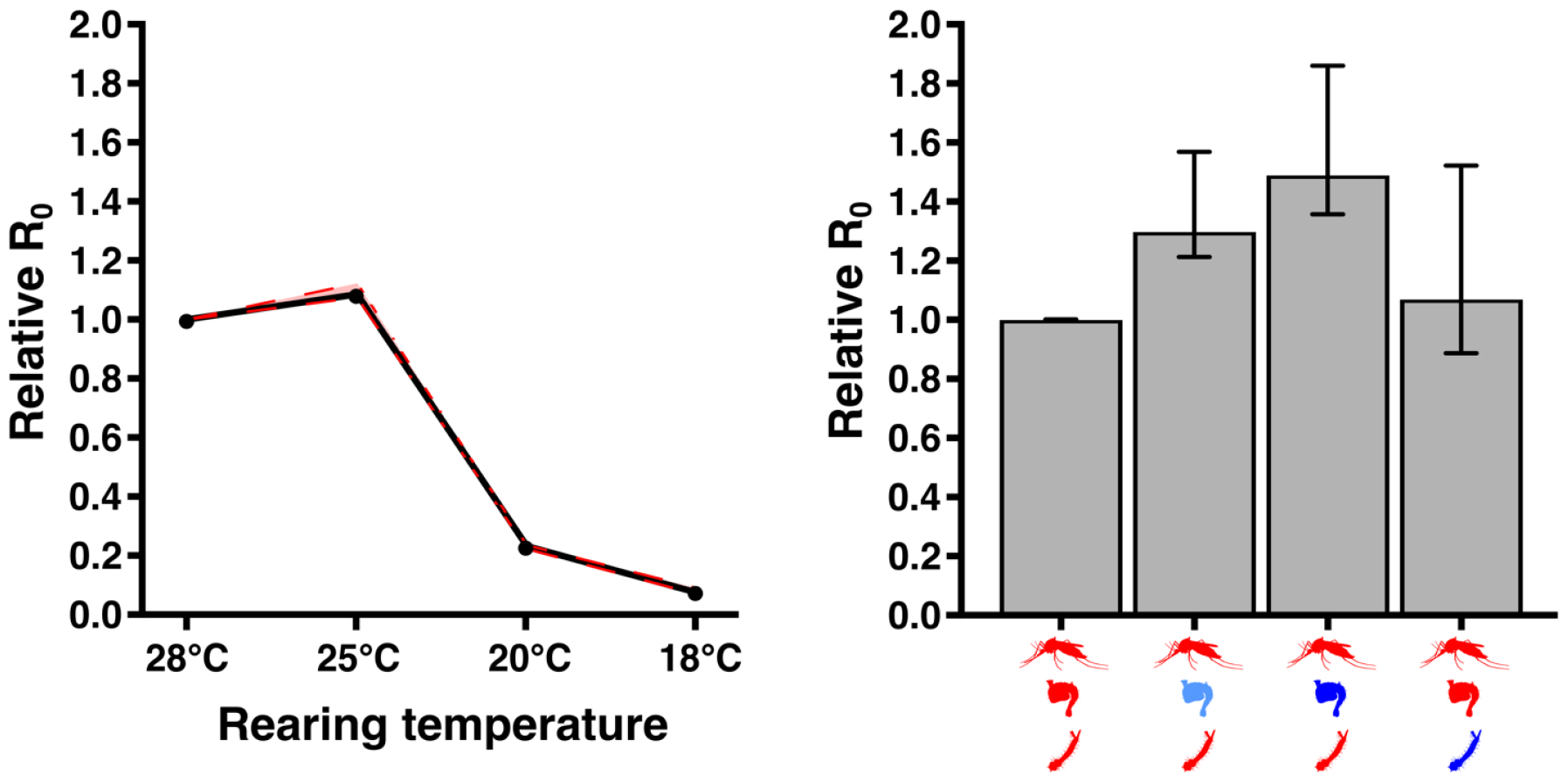
Temperature-dependent modeling of relative ZIKV *R*_0_ values. **(A)** Relative *R*_0_ of ZIKV when vectored by *Ae. aegypti* reared at a single constant low temperature before virus exposure, in contrast to the control maintained at a constant 28°C. The 95% confidence interval is indicated by a red dashed line. (**B**) Relative *R*_0_ of ZIKV when vectored by *Ae. aegypti* exposed to colder temperatures during either the pupal or larval stages, relative to the control held at a constant 28°C. Error bars indicate 95% confidence interval. Symbol colors denote the temperature at which each life stage was maintained, with red indicating 28°C, light blue indicating 25°C, and dark blue indicating 18°C.

## Discussion

Exposure to cooler temperatures during the development of mosquitoes has been suggested to reduce barriers to midgut infection and escape, thereby enhancing vector competence for arbovirus pathogens (18, 32). However, many laboratory-based studies exploring temperature-dependent effects on vector competence, as well as mechanistic models incorporating environmental conditions, often overlook the natural thermal environments of disease vectors (4, 14, 28, 33). Instead, a significant portion of research on mosquito vector competence is conducted under constant temperatures, which do not mirror the diurnal temperature fluctuations experienced by mosquitoes in their natural habitats (21, 25). Studies that do investigate the impact of temperature on vector competence usually focus solely on the extrinsic incubation period (EIP) of the pathogen. Notably, Tesla et al (28) found the impact of temperature on *Ae. aegypti* following ZIKV ingestion to be non-linear, revealing a unimodal relationship. This study estimated the optimal transmission temperature to be 30.6°C. However, the temperature range conducive to virus transmission was delimited by minimum and maximum temperatures of 22.9°C and 38.4°C, respectively. The transmission of ZIKV seems to be restricted by distinct mechanisms at these thermal extremes. While higher temperatures rendered the mosquitoes more susceptible to infection, increases in mosquito mortality constrained overall transmission potential. Lower temperatures appeared to limit the overall numbers of infectious mosquitoes by affecting the physiology of the mosquito and influencing the dynamics of the virus infection itself (28). Similar temperature dependent effects during the EIP have also been observed in other virus-vector systems (6, 21). While these studies are important for understanding the effects of temperature following ingestion of the virus, our results suggest that pathogen transmission is also influenced by the heterogenous temperature conditions occurring prior to virus introduction, during the immature stages of mosquito development; in particular, the pupal phase.

Mathematical models used to calculate vectorial capacity often assume a consistent temperature throughout the mosquito’s life cycle. However, this assumption overlooks the fact that aquatic immature stages and terrestrial adults may encounter different environmental conditions in habitats with diverse microclimates (34–36). Notably, our group and others have demonstrated that exposure to cooler temperatures solely during the juvenile developmental stages increases the susceptibility of adult mosquitoes to arbovirus infections (17–21). Mosquitoes are ectothermic, meaning that internal body temperatures are not regulated and fluctuate with the thermal conditions of the surrounding environment. Consequently, environmental temperatures play a crucial role not only in pathogen-host dynamics but also in influencing various other processes that impact the biology of disease vectors, such as development time (10, 11, 37) and behaviors (38–41). Previous modeling studies have anticipated a decrease in arbovirus transmission by mosquitoes in colder climates due to prolonged development times, reduced biting rates, and an extended EIP (4, 28, 29, 42, 43). However, the findings presented in this study reveal that subjecting only the pupal stage of *Ae. aegypti* to fluctuating diurnal temperatures < 23°C significantly increases the efficiency of ZIKV transmission (Fig. 4E). While mosquitoes reared at either constant low temperatures or fluctuating diurnal temperatures were equally susceptible to ZIKV, the transmission rate increased significantly when mosquitoes were also exposed to higher fluctuating temperatures during the EIP (Fig. 4E; regime IV). Simulations run with these parameters, with pupae subjected to temperatures ≤ 25°C, predict higher *R*_0_ values for ZIKV (Fig. 5B). This phenomenon is likely attributable to the relatively brief duration of the pupal stage at these lower temperatures (≤ 4 days). Exposure to cooler temperatures during this life stage significantly increased vector competence, with minimal impact on overall development time. An extended development time typically correlates with reduced survival (10, 37). For instance, the relative *R*_0_ value for ZIKV is predicted to decrease when all immature life stages and early adult *Ae. aegypti* are reared at 18°C prior to virus exposure (Fig. 5A). This is likely because the observed increases in vector competence at this temperature (Fig. 2D) are counteracted by an overall decrease in mosquito longevity, resulting from a significantly extended development time at this temperature (6, 10, 37, 44). These results contrast with the higher relative *R*_0_ value computed for ZIKV if only the pupal stage is exposed to 18°C prior to virus exposure (Fig. 5B).

In summary, the data presented here emphasizes the importance of considering the environmental conditions that vector species encounter throughout their life cycle when evaluating the risk posed by mosquito-borne viruses. Although it is impractical to test every conceivable environmental condition found in nature, shortcomings in the experimental design of many studies on vector competence, coupled with flawed assumptions in current mathematical models, may be impeding our ability to predict the spread of diseases transmitted by mosquitoes. In particular, the environmental conditions experienced during the pupal stage of *Ae. aegypti* development may play a more important role in the transmission dynamics of ZIKV than previously recognized. To enhance the accuracy of predictive models for mosquito-borne viruses, future research should explore the physiological basis of the phenotypes described here, and adopt more comprehensive experimental designs that consider diurnal temperature fluctuations. Previous studies have indicated that temperature-dependent effects on innate immune pathways could impact the transmission dynamics of mosquito-borne pathogens (7, 24, 45–48). For example, our laboratory previously observed inhibition of the primary antiviral pathway in *Ae. aegypti* after rearing at 18°C, which correlated with an increased susceptibility to arboviral infections. This offers a potential molecular mechanism for observed temperature-dependent changes in vector competence (17). Although our current study did not delve into this possibility or investigate potential physiological explanations, further research in this area could provide valuable insights into the transmission of significant viral diseases by mosquitoes.

## Materials and Methods

### Experimental Design

We utilized programmable environmental chambers capable of ramping and soaking (Caron, Marietta, Ohio, USA) to determine the effects of low constant and fluctuating temperatures prior to virus exposure on the vector competence of *Ae. aegypti* for ZIKV. In the first set of experiments, mosquitoes were reared at different constant temperatures that were lower than the usual insectary conditions for *Ae. aegypti* (18°C, 23°C, or 25°C). Second, we exposed only the pupal or larval stages to 18°C or 25°C while holding other stages at 28°C. Mosquitoes reared at a constant 28°C prior to introduction of ZIKV were considered the control for these experiments. Lastly, we worked with Harris County Mosquito Control to deploy data loggers (HOBO Pendant MX Temperature/Light data logger, Onset, Bourne, MA, USA) in mosquito microhabitats near traps sites across Harris County, TX. Temperatures were recorded at 10-minute intervals over a period of one year. Using the data collected, we simulated five different fluctuating temperature regimes corresponding with the diurnal temperature ranges recorded at each site from September to November when arbovirus transmission is historically higher than other times of years. The parameters of each fluctuating diurnal temperature regimes were as follows: regime I had a of DTR of 7.8°C around a mean of 29°C (min: 25.1°C, max: 32.9°C) prior to blood feeding and moved to a constant 28°C after virus exposure; regimes II and III had a DTR of 7.8°C around a mean of 19°C (min: 15.1°C, max :22.9°C) during only the pupal or pupal and adult stages, respectively, but otherwise followed regime I; regime IV was the same as regime III but returned to a DTR of 7.8°C around a mean of 29°C (min: 25.1°C, max: 32.9°C) after virus exposure; regime V had a DTR of 3.5°C around a mean of 19°C (min: 17.3°C, max: 20.3°C) and moved to a constant 28°C after virus exposure. All mosquitoes were reared and maintained under a 14:10hr day:night cycle with 70-80% relative humidity. Temperatures and relative humidity in the environmental chambers were monitored and recorded during all experiments using HOBO data loggers (Onset, Bourne, MA, USA).

### Mosquitoes

*Ae. aegypti* (Liverpool strain) were utilized for all experiments. Colony maintenance was performed under standard insectary conditions of a constant 28°C with 70-80% humidity and a 14:10hr day:night cycle.

Mosquito larvae were fed fish food and adults were maintained on 10% sucrose solution.

### Experimental infections

ZIKV (PRVABC059) was rescued from an infectious clone and passaged two times on Vero E6 cells (ATCC, CRL-1586) and three times on C6/36 cells (ATCC, CRL-1660) cultured in DMEM supplemented with 10% heat-inactivated FBS and 1% penicillin-streptomycin at 37°C and 28°C w/ 5% CO_2_, respectively. ZIKV stocks were prepared and frozen at −80°C. Virus stock titrations were done by plaque assay on Vero E6 cells. For each biological replicate, ∼100 three-to-five-day old *Ae. aegypti* were offered a blood meal containing 1 x 10^6^ PFU/ml ZIKV heated to 37°C using a Hemotek feeding apparatus. Blood meals were prepared by mixing defibrinated sheep’s blood (Colorado Serum Company) in a 1:1 ratio with virus suspension. Mosquitoes were allowed to feed for 1hr then fully engorged females were separated into individual chambers with 30-45 mosquitoes per biological replicate. Eight hours prior to blood feeding, all mosquitoes were moved to 28°C for 8hr to reduce negative effects that lower temperatures may have on host seeking behavior (6, 38, 39). Three full biological replicates were performed for each set of experiments.

### Vector competence

Mosquitoes were assessed for ZIKV midgut infection, systemic infection, and transmission 14 days post blood meal. Prevalence of ZIKV midgut and systemic infections was determined by cytopathic effects (CPE) assay on BHK-21 cells (ATCC, CCL-10) as previously described (49), and viral load was quantified by strand-specific quantitative real-time PCR (RT-qPCR). Transmission of ZIKV by *Ae. aegypti* was assessed using a modified honey-card method (50–53). Briefly, mosquitoes were placed into individual chambers following blood feeding and maintained on filter papers (Whatman, Piscataway, New Jersey) soaked in 10% sucrose solution. The filter papers can be examined for the presence of ZIKV genome as the mosquitoes expectorate on the filter papers during feeding (rate of transmission of ZIKV [# positive filter papers / # positive heads], transmission efficiency [# positive filter papers / # exposed]). Viral RNA was extracted from individual mosquito bodies and filter papers using the Nucleospin RNA Virus kit (Machery-Nagel, Düren, Germany) according to the manufacturer’s protocol. ZIKV genomic strands were detected and quantified by strand-specific quantitative real-time PCR (RT-qPCR) as previously described (14). The specific primers and probe used were as follows: ZIKV F, 5’-TGCCCATACACCAGCACTATG-3’, ZIKV R, 5’-GGCCGTCATGGTGGCGAATAA-3’, ZIKV probe, 5’-CCTGGAGCGACTGCAGCGTAG GTATGGGGG-3’.

### Modeling approach

We extended an existing temperature-dependent Zika *R*_0_ model derived from a modified version of the Ross-MacDonald model for mosquito-borne virus transmission (4, 28, 31). This widely used *R*_0_ model is defined as :

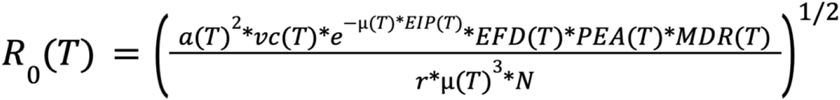

with the parameters for this temperature-dependent model given as follows: *a* is the per-mosquito biting rate; *vc* is vector competence; *μ* is the mosquito mortality rate; *EIP* is the extrinsic incubation period; *EFD* is the number of eggs produced per female mosquito per day; *MDR* is the mosquito development rate; *p*_*EA*_ is the probability of mosquitoes surviving to adulthood; *N* is the density of human hosts; and *r* is the recovery of rate of human hosts. *MDR* is equal to 1/(t_*E*_ + t_*L*_ + t_*P*_) where t_*i*_ is the duration of time as eggs (E), larvae (L), and pupae stage (P), respectively. With *T* being the temperature in degrees celsius.

We extended this *R*_0_ model by differentiating the role of aquatic stage temperature, *T*_0_, from adult stage temperature, *T*, on *R*_0_, and define the new model as follows:

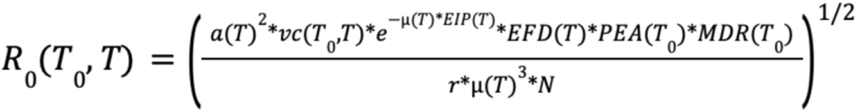

With the exception of the vector competency, *vc*, whose values were obtained from our laboratory experiments, the temperature-dependent parameter functions t_*i*_(*T*_*0*_) were obtained from Erraguntla et al. (54), and the other parameters were obtained from previous data-driven modeling studies (4, 28, 55). To evaluate the impact of *Ae. aegypti* aquatic stages temperature on Zika transmission risk, we computed the relative *R*_0_ value as the ratio of *R*_0_(*T*_0_,*T*) for a given aquatic and adult stage temperature, *T*_0_ and *T*, respectively, over *R*_0_ (28,28), with 28°C. This temperature was identified by previous modeling study as near-optimal for ZIKV transmission by *Ae. aegypti* (28).

### Statistical analysis

All data were analyzed using GraphPad Prism 9 software. Susceptibility and transmission data were assessed by performing a one-way ANOVA and Dunnett’s multiple comparisons test on three independent biological replicates. Quantification of ZIKV genomes by RT-qPCR was assessed by performing a one-way ANOVA and Dunnett’s multiple comparisons test on three independent biological replicates with three technical machine replicates.

## Acknowledgments

We thank Fang Wang, Brooke Norwood, and Victoria Wong for assistance in rearing mosquitoes.

## Author Contributions

T.D.P, Z.N.A, and K.M.M designed research; T.D.P, B.H., M.L, M.E, M.R, M.D, C.F., J.V., and K.M.M. performed research; M.N. implemented the temperature dependent model; T.D.P, M.N., and K.M.M analyzed data; T.D.P and K.M.M wrote the manuscript.

